# Identification of a novel cationic glycolipid in *Streptococcus agalactiae* that contributes to brain entry and meningitis

**DOI:** 10.1101/2020.12.01.406702

**Authors:** Luke R. Joyce, Haider S. Manzer, Jéssica da C. Mendonça, Ricardo Villarreal, Prescilla E. Nagao, Kelly S. Doran, Kelli L. Palmer, Ziqiang Guan

## Abstract

Bacterial membrane lipids are critical for membrane bilayer formation, cell division, protein localization, stress responses, and pathogenesis. Despite their critical roles, membrane lipids have not been fully elucidated for many pathogens. Here, we report the discovery of a novel cationic glycolipid, Lysyl-Glucosyl-Diacylglycerol (Lys-Glc-DAG) that is synthesized in high abundance by the bacterium *Streptococcus agalactiae* (Group B *Streptococcus*, GBS). To our knowledge, Lys-Glc-DAG is more positively charged than any other known lipids. Lys-Glc-DAG carries two positive net charges per molecule, distinct from the widely described lysylated phospholipid Lysyl-phosphatidylglycerol (Lys-PG) which carries one positive net charge due to the presence of a negatively charged phosphate moiety. We use normal phase liquid chromatography (NPLC) coupled with electrospray ionization (ESI) high-resolution tandem mass spectrometry (HRMS/MS) and genetic approaches to determine that Lys-Glc-DAG is synthesized by the enzyme MprF in GBS, which covalently modifies the neutral glycolipid Glc-DAG with the cationic amino acid lysine. GBS is a leading cause of neonatal meningitis, which requires traversal of the endothelial blood-brain barrier (BBB). We demonstrate that GBS strains lacking *mprF* exhibit a significant decrease in the ability to invade BBB endothelial cells. Further, mice challenged with a GBSΔ*mprF* mutant developed bacteremia comparably to Wild-Type infected mice yet had less recovered bacteria from brain tissue and a lower incidence of meningitis. Thus, our data suggest that Lys-Glc-DAG may contribute to bacterial uptake into host cells and disease progression. Importantly, our discovery provides a platform for further study of cationic lipids at the host-pathogen interface.

## Introduction

Bacterial cellular membranes are dynamic structures that are critical for survival under varying environmental conditions and are essential for host-pathogen interactions. Phospholipids and glycolipids within the membrane have varying chemical properties that alter the physiology of the membrane, which bacteria can modulate in response to environmental stresses such as pH (1), antibiotic treatment (2), and human metabolites (3). Despite their critical roles in the survival and pathogenesis, membrane lipids have not been carefully characterized using modern lipidomic techniques for many important human pathogens, including *Streptococcus agalactiae* (Group B *Streptococcus*; GBS). GBS colonizes the lower genital and gastrointestinal tracts of ∼30% of healthy women (4, 5). However, GBS can cause sepsis and pneumonia in newborns and is a leading cause of neonatal meningitis, resulting in long-lasting neurological effects in survivors (6-8). Due to the severity of the resulting diseases, intrapartum antibiotic prophylaxis is prescribed for colonized pregnant women (7, 9). Even with these measures, a more complete understanding of GBS pathogenesis and new therapeutic and preventive measures are needed to mitigate the devastating impact of GBS neonatal infection.

Research on the pathogenesis of the GBS has mainly focused on cell wall-anchored or secreted proteins and polysaccharides that aid in the attachment to and invasion of host cells. The numerous attachment and virulence factors possessed by the GBS are summarized in a recent review by Armistead *et al* 2019 (10). Comparatively little is known about GBS cellular membrane lipids. To our knowledge, the only characterization of GBS lipids prior to our current study was the identification of the phospholipids phosphatidylglycerol (PG), cardiolipin (CL), and lysyl-phosphatidylglycerol (Lys-PG) in GBS (11-13). Similarly, investigation into the glycolipids of the GBS membrane has focused on di-glucosyl-diacylglycerol (Glc2-DAG), which is the lipid anchor of the Type I lipoteichoic acid, and its role in pathogenesis (14).

In this study, we utilized normal phase liquid chromatography (NPLC) coupled with electrospray ionization (ESI) high-resolution tandem mass spectrometry (HRMS/MS) to characterize the GBS membrane lipid composition, and identified a novel cationic glycolipid, lysyl-glucosyl-diacylglycerol (Lys-Glc-DAG), which comprises a major portion of the GBS total lipid extract. While Lys-PG has been reported in a range of bacterial species (15), Lys-Glc-DAG represents, to our knowledge, the first example of lysine modification of a neutral glycolipid. By gene deletion and heterologous expression, we show the GBS MprF enzyme is responsible for the biosynthesis of both the novel Lys-Glc-DAG and Lys-PG. Most strikingly, using an *in vivo* hematogenous murine infection model, we demonstrate that MprF does not contribute to GBS bloodstream survival. This distinguishes the GBS MprF from the well-known *Staphylococcus aureus* MprF, which synthesizes only Lys-PG (16, 17). Rather, GBS MprF contributes specifically to meningitis and penetration of the blood-brain barrier. These results greatly expand our knowledge of naturally occurring lipids and MprF functionality and reveal insights into the pathogenesis of meningitis caused by GBS.

## Results

### Identification of Lys-Glc-DAG, a novel cationic glycolipid in GBS

The membrane lipids of three GBS clinical isolates of representative serotypes were characterized: COH1 (18), A909 (19), CNCTC 10/84 and CJB111 (20) (serotypes III, 1a, and V, respectively). Common Gram-positive bacterial lipids were identified by normal phase LC coupled with negative ion ESI/MS/MS, including diacylglycerol (DAG), monohexosyldiacylglycerol (MHDAG), dihexosyldiacylglycerol (DHDAG), phosphatidylglycerol (PG), and lysyl-phosphatidylglycerol (Lys-PG), as shown by the negative total ion chromatogram (TIC) (Fig. 1A).

**Fig 1.**
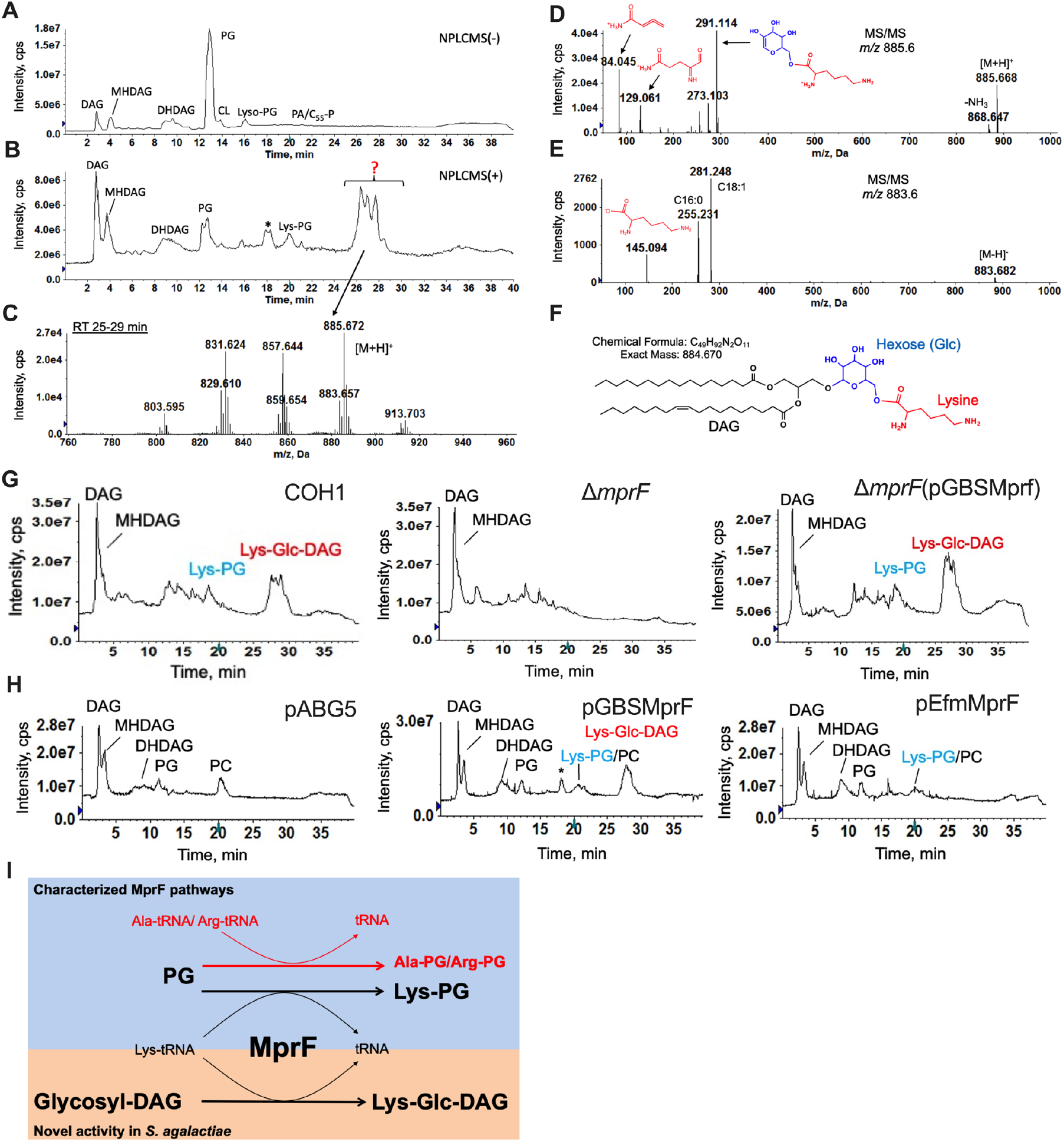
Lipidomic profiling of GBS and identification of Lys-Glc-DAG synthesized by MprF. Total ion chromatogram (TIC) of LC/MS in A) negative ion mode, B) positive ion mode shows a major unknown lipid eluting at ∼25-29 min. C) Positive ESI/MS showing the [M+H]^+^ ions of the unknown lipid. D) Positive ion MS/MS spectrum of [M+H]^+^ at *m/z* 885.6 and E) negative ion MS/MS spectrum of [M-H]^-^ at *m/z* 883.6 of the unknown lipid. F) Lys-Glc-DAG (16:0/18:1) is proposed as the structure of the unknown lipid. G) TIC showing loss of Lys-Glc-DAG and Lys-PG in COH1Δ*mprF* which is present when *mprF* is complemented *in trans*. H) Lys-Glc-DAG and Lys-PG is only present in *S. mitis* when expressing GBS *mprF* compared to Lys-PG only when expressing *E. faecium mprF*. “*” denotes methylcarbamate of Lys-Glc-DAG, an extraction artifact due to the use of chloroform. I) Biosynthetic pathways involving MprF.

Surprisingly, the positive TIC (Fig. 1B, S1 Fig) shows highly abundant peaks of unknown identity at the retention time ∼25-29 min. The mass spectra (Fig. 1C) and LC retention times of this lipid do not match with any other bacterial lipids we have analyzed or exact masses in lipidomic databases (21, 22). Tandem MS (MS/MS) in the positive ion mode (Fig. 1D), negative ion mode (Fig. 1E), and high-resolution mass measurement (Fig. 1C) allowed us to propose lysyl-glucosyl-diacylglycerol (Lys-Glc-DAG) (Fig. 1F) as the structure of this unknown lipid. Observed and exact masses of Lys-Glc-DAG are shown in S1 Table. The assignment of glucose was based on the observation that glucosyl-diacylglycerol (Glc-DAG) is a major membrane component of GBS and other streptococci (14), and results from an isotopic labeling experiment using ^13^C-labeled glucose (Glucose-^13^C6). The assignment of lysine modification was supported by an isotopic labeling experiment with deuterated lysine (lysine-*d4*). The expected mass shifts (+4 Da) were observed in both molecular ions and MS/MS product ions (S2 Fig). Comparison of both MS/MS spectra of labeled (Glucose-^13^C_6_) and unlabeled Lys-Glc-DAG indicates the lysine residue is linked to the 6-position of glucose (S2 Fig). Lys-Glc-DAG consists of several molecular species with different fatty acyl compositions resulting in different retention times and multiple, unresolved TIC peaks (∼25-29 min).

### GBS MprF synthesizes Lys-Glc-DAG

The enzyme MprF (multiple peptide resistance factor) catalyzes the aminoacylation of PG with lysine in some Gram-positive pathogens (16, 23). We determined that GBS MprF is responsible and sufficient for synthesizing Lys-Glc-DAG as well as Lys-PG. Deletion of *mprF* from both COH1 and CJB111 abolishes Lys-Glc-DAG and Lys-PG synthesis, which are restored by complementation (Fig. 1G, S3 Fig). Deletion of GBS *mprF* does not confer a growth defect in Todd-Hewitt broth or tissue culture medium. The oral colonizer *Streptococcus mitis* does not encode *mprF* or synthesize Lys-PG but synthesizes Glc-DAG and PG (2, 3). Heterologous expression of GBS *mprF* in *S. mitis* results in Lys-Glc-DAG and Lys-PG production (Fig. 1H), while expression of *Enterococcus faecium mprF* results in only Lys-PG production (Fig. 1H), as expected (1). Biosynthetic pathways involving MprF are shown in Fig. 1I.

### MprF contributes to GBS pathogenesis

We investigated whether MprF contributes to GBS invasion into brain endothelial cells and development of meningitis. To mimic the human blood-brain barrier (BBB), we utilized the human cerebral microvascular endothelial cell line hCMEC/D3. *In vitro* assays for adhesion and invasion were performed as described previously (14, 24, 25). There was no significant difference in the ability of Δ*mprF* compared to WT and complement cells to attach to hCMEC/D3 cells (Fig. 2A). However, we observed a significant decrease in the amount of Δ*mprF* recovered from the intracellular compartment of hCMEC/D3 cells (Fig. 2A). The reduced invasion phenotype was confirmed in the hypervirulent serotype V strain, CJB111 (26, 27) (S4 Fig). Intracellular survival requires GBS to survive low pH conditions in lysosomes (pH 4.5 – 5.5) (28), and *ΔmprF* is unable to survive low pH conditions (Fig. 2B). This suggests that MprF promotes GBS invasion and possibly intracellular survival in brain endothelial cells.

**Fig 2.**
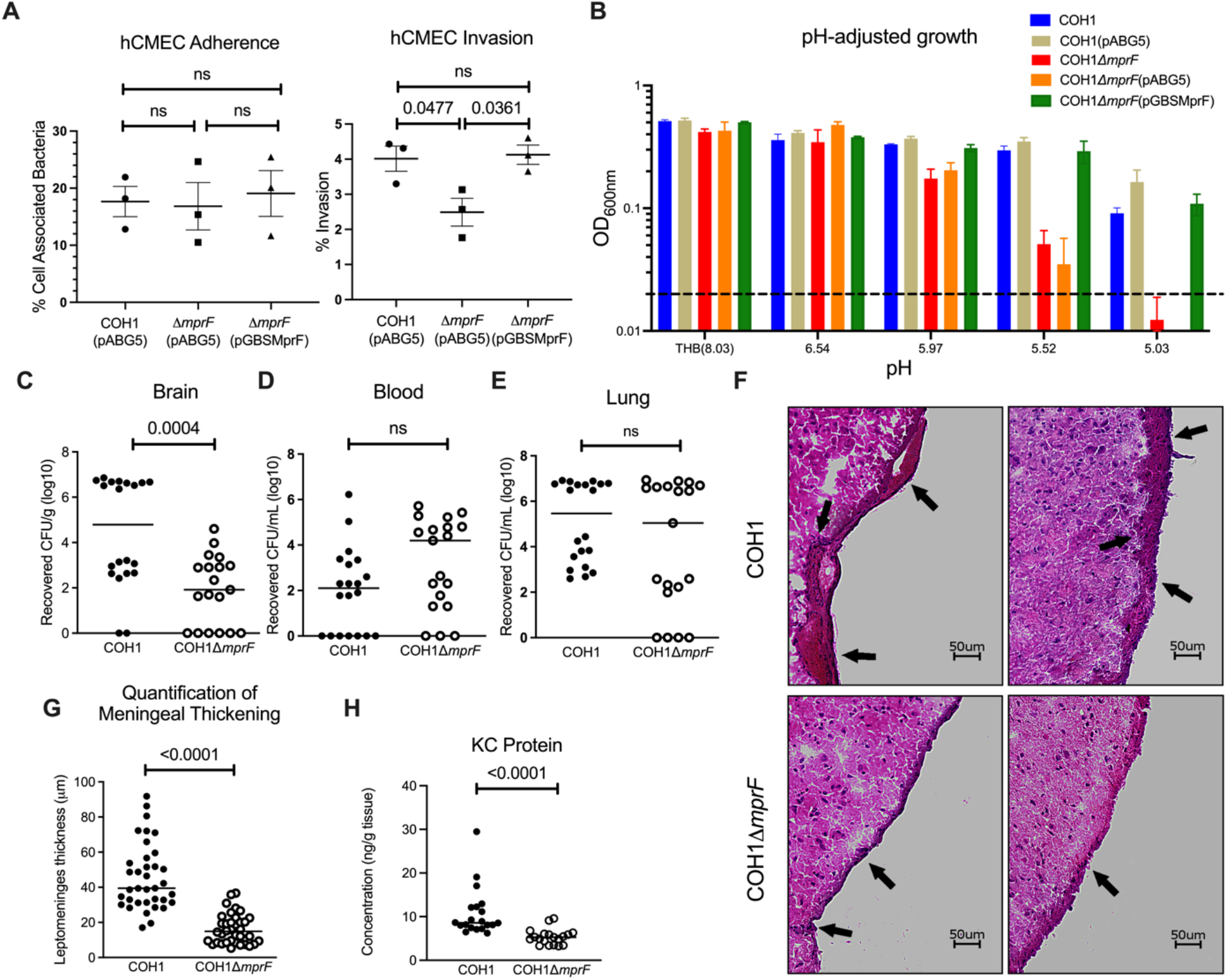
Contribution of lysine lipids to meningitis pathogenesis. A) *In vitro* assays for adherence and invasion of hCMEC cells indicates *mprF* contributes to invasion but not adherence to brain endothelium (the mean of each biological replicate is displayed, comprised of 4 replicate wells per biological replicate, mean and SEM). B) pH-adjusted medium growth indicates *ΔmprF* cannot survive in low pH conditions, mean and SD. Groups of CD-1 mice were injected intravenously with COH1 WT or COH1Δ*mprF* strains and bacterial counts were assessed in the C) brain, D) blood, and E) lung after 72h. Representative data from 2 independent experiments are shown (WT, *n =* 20; Δ*mprF, n* = 19). F) Hematoxylin-eosin-stained brain sections from representative mice infected with WT (top) or Δ*mprF* mutant (bottom); arrows indicate meningeal thickening and leukocyte infiltration. G) Quantification of meningeal thickening using ImageJ. H) KC chemokine production measured by ELISA. Panels C, D, E, G, and H) median indicated. Statistical analyses performed using GraphPad Prism: A) One-way ANOVA with Tukey’s multiple comparisons test; C, D, F) unpaired two-tailed t-test; E, G) Mann-Whitney U test; p-values indicated; ns, no significance (p-value > 0.05).

We hypothesized that these *in vitro* phenotypes of Δ*mprF* would translate into a diminished ability to penetrate the BBB and produce meningitis *in vivo*. Using our standard model of GBS hematogenous meningitis (14, 24) mice were challenged with either WT GBS or Δ*mprF*. Mice were sacrificed at 72 h to determine bacterial loads in blood, lung and brain tissue. We recovered significantly less CFU in the brains of Δ*mprF*-infected mice compared to the WT-infected mice (Fig. 2C). However, there was no significant difference in CFU recovered from the bloodstream or the lung (Fig. 2D,E), demonstrating that Δ*mprF* does not have a general defect in bloodstream survival or tissue invasion *in vivo*. Furthermore, mice challenged with WT GBS had significantly more meningeal thickening and neutrophil chemokine, KC, in brain homogenates compared to Δ*mprF* mutant-infected animals (Fig. 2F-H). Taken together, *mprF* contributes to GBS penetration into the brain and to the pathogenesis of meningitis *in vivo*.

## Discussion

Here, we report that GBS MprF uniquely synthesizes a novel cationic glycolipid Lys-Glc-DAG in high abundance plays a role in the invasion of human endothelial cells. This work establishes that GBS capitalizes on MprF to modulate charges of both glycolipids and phospholipids at the membrane, which is unprecedented

Previously MprF has been shown to catalyze the aminoacylation of the anionic phospholipid PG in a range of Gram-positive and Gram-negative bacteria (16, 23). MprF is a membrane-bound enzyme comprised of a N-terminal lipid flippase domain (29) and a C-terminal catalytic domain that catalyzes the aminoacylation of the glycerol group of PG by using aminoacyl-tRNAs as the amino acid donors (30-32). An important function of PG aminoacylation is proposed to decrease the net negative charge of the cellular envelope to confer protection from cationic antimicrobial peptides (CAMPs) produced by host immune systems and bacteriocins produced by competitor bacteria (16, 23). However, a previous study observed no contribution of *mprF* to GBS *in vitro* susceptibility to commonly studied CAMPs, which is unlike the well-characterized *S. aureus mprF* (33), thus highlighting the unique differences between the extracellular surface of these bacteria.

Based on our tissue culture and mouse infection experiments, we propose that GBS have an MprF enzyme and corresponding cellular lipid properties that are adapted for efficient invasion of mammalian cells. Deletion of *mprF* impacts the ability of GBS to enter the brain and promote meningitis *in vivo*. This suggests that MprF plays a role in BBB penetration and not invasion into the lung, however additional studies are warranted to examine other tissue sites. It is unknown how lysinylated lipids in the GBS membrane, which is covered by a layer of peptidoglycan, mechanistically impact invasion. Because Lys-Glc-DAG is abundantly synthesized by GBS MprF, with Lys-PG a comparatively minor product, it is likely that Lys-Glc-DAG is the most relevant lipid for meningitis pathogenesis. Speculatively, Lys-Glc-DAG may contribute to membrane vesicle (MV) formation by GBS. MVs have previously been shown to be pro-inflammatory and result in preterm birth and fetal death in mice (34), but have not been studied during meningitis progression. In future studies, it will be key to investigate this, as well as the specific host inflammatory and signaling responses to the GBS *mprF* mutant.

Our identification of the novel Lys-Glc-DAG glycolipid rationalizes further study of the lipidomes of human pathogens. First, lipids contribute to virulence, and understanding these virulence mechanisms and the mechanisms for lipid synthesis may identify novel antimicrobial drug targets. The decreased *in vivo* pathogenicity of the Δ*mprF* mutant identifies GBS MprF as a candidate for targeting by antimicrobial strategies. Moreover, Lys-Glc-DAG could be utilized as a specific molecular biomarker for GBS diagnostics. In addition, engineered cationic lipids are utilized in lipid nanoparticles for mRNA vaccine and drug delivery and are required for uptake of particles into cells (35, 36). Substantial effort has been dedicated to the synthesis of cationic lipids with low toxicity and efficient delivery properties. Lys-Glc-DAG is a naturally occurring, strongly cationic lipid with potential for use in lipid nanoparticles for vaccine and drug delivery. Importantly, our discovery suggests that lipidome analysis of human pathogens is likely to reveal novel lipids of biotechnological utility.

## Materials and methods

### Bacterial strains, media, and growth conditions

See S2 Table for strains used in this study. GBS strains were grown statically at 37°C in Todd-Hewitt Broth (THB) and *S. mitis* strains were grown statically at 37°C and 5% CO_2_, unless otherwise stated. Streptococcal chemically defined medium (37) was diluted from stock as described (38) with 1% w/v glucose (referred to as DM), slightly modified from (39), unless otherwise stated. *Escherichia coli* strains were grown in Lysogeny Broth (LB) at 37°C with rotation at 225 rpm. Kanamycin and erythromycin (Sigma-Aldrich) were supplemented to media at 50 µg/mL and 300 µg/mL for *E. coli*, respectively, or 300 µg/mL and 5 µg/mL, respectively, for streptococcal strains.

### Routine molecular biology techniques

All PCR reactions utilized Phusion polymerase (Thermo Fisher). PCR products and restriction digest products were purified using GeneJET PCR purification kit (Thermo Fisher) per manufacturer protocols. See S3 Table for primers. Plasmids were extracted using GeneJET plasmid miniprep kit (Thermo Fisher) per manufacturer protocols. Restriction enzyme digests utilized XbaI, XhoI, and PstI (New England Biolabs) for 3 h at 37°C in a water bath. Ligations utilized T4 DNA ligase (New England Biolabs) at 16°C overnight or Gibson Assembly Master Mix (New England Biolabs) per manufacturer protocols where stated. All plasmid constructs were sequence confirmed by Sanger sequencing (Massachusetts General Hospital DNA Core or CU Anschutz Molecular Biology Core).

### Deuterated lysine and ^13^C_6_-D-glucose isotope tracking

A GBS COH1 colony was inoculated into 15 mL of DM containing 450 µM lysine-*d4* (Cambridge Isotopes Laboratories) or a single COH1 colony was inoculated into 10 mL DM supplemented with 0.5% w/v ^13^C_6_D-glucose (U-13C6, Cambridge Isotopes Laboratories) for overnight growth for lipidomic analysis described below.

### Construction of MprF expression plasmids

Genomic DNA was isolated using the Qiagen DNeasy Blood and Tissue kit per the manufacturer’s protocol with the exception that cells were pre-treated with 180 µL 50 mg/mL lysozyme, 25 µL 2500 U/mL mutanolysin, and 15 µL 20 mg/mL pre-boiled RNase A and incubated at 37°C for 2 h. The *mprF* genes from GBS COH1, (GBSCOH1_1931), GBS CJB111 (ID870_10050), and *E. faecium* 1,231,410 (EFTG_00601) were amplified and either Gibson ligated into pABG5Δ*phoZ* (40) or ligated into pDCErm (41). Plasmid constructs were transformed into chemical competent *E. coli*. Briefly, chemically competent cells were incubated for 10 min on ice with 5 µL of Gibson reaction before heat shock at 42°C for 70 sec, then placed on ice for 2 min before 900 µL of cold SOC medium was added. Outgrowth was performed at 37°C, with shaking at 225 rpm, for 1 h. Cultures were plated on LB agar plates containing 50 µg/mL kanamycin. Colonies were screened by PCR for presence of the *mprF* insert.

### Expression of *mprF* in *S. mitis*

Natural transformation was performed as previously described (3). Briefly, precultures were thawed at room temperature, diluted in 900 µL of THB, further diluted 1:50 in prewarmed 5 mL THB, and incubated for 45 min at 37°C. 500 µL of culture was then aliquoted with 1 µL of 1 mg/ml competence-stimulating peptide (EIRQTHNIFFNFFKRR) and 1 µg/mL plasmid. Transformation reaction mixtures were cultured for 2 h at 37°C in microcentrifuge tubes before being plated on THB agar supplemented with 300 µg/mL kanamycin. Single transformant colonies were cultured in 15 mL THB overnight. PCR was used to confirm the presence of the *mprF* insert on the plasmid. Plasmids were extracted and sequence confirmed as described above. Lipidomics was performed as described below in biological triplicate.

### Construction of *mprF* deletion plasmids

Regions ∼2 kb upstream and downstream of the GBS COH1 *mprF* (GBSCOH1_1931) or CJB111 (ID870_10050) were amplified using PCR. Plasmid, pMBSacB (42), and the PCR products were digested using appropriate restriction enzymes and ligated overnight. 7 µL of the ligation reaction was transformed into chemically competent *E. coli* DH5α as described above, except that outgrowth was performed at 28°C with shaking at 225 rpm for 90 min prior to plating on LB agar supplemented with 300 µg/mL erythromycin. Plates were incubated at 28°C for 72 h. Colonies were screened by PCR for correct plasmid construction. Positive colonies were inoculated into 50 mL LB media containing antibiotic and incubated at 28°C with rotation at 225 rpm for 72 h. Cultures were pelleted using a Sorvall RC6+ centrifuge at 4,280 x *g* for 6 min at room temperature. Plasmid was extracted as described above except the cell pellet was split into 5 columns to prevent overloading and serial eluted into 50 µL. Plasmid construction was confirmed via restriction digest using XhoI and XbaI, and the insert was PCR amplified and sequence-verified.

### Generation of electrocompetent GBS cells for *mprF* knockout

Electrocompetent cells were generated as described (42) with minor modifications. Briefly, a GBS COH1 or CJB111 colony was inoculated in 5 mL M17 medium (BD Bacto) with 0.5% glucose and grown overnight at 37°C. The 5 mL was used to inoculate a second overnight culture of 50 mL pre-warmed filter-sterilized M17 medium containing 0.5% glucose, 0.6% glycine, and 25% PEG 8000. The second overnight was added to 130 mL of the same medium and grown for 1 h at 37°C. Cells were pelleted at 3,200 x *g* in a Sorvall RC6+ at 4°C for 10 min. Cells were washed twice with 25 mL cold filter-sterilized GBS wash buffer containing 25% PEG 8000 and 10% glycerol in water, and pelleted as above. Cell pellets were re-suspended in 1 mL GBS wash buffer and either used immediately for transformation or stored in 100 µL aliquots at -80°C until use.

### Deletion of GBS COH1 and CJB111 *mprF*

Electrocompetent cells were generated as described (42) with minor modifications. The double crossover homologous recombination knockout strategy was performed as described previously (25, 42, 43) with minor modifications. 1 µg of plasmid was added to electrocompetent GBS cells and transferred to a cold 1 mm cuvette (Fisher or BioRad). Electroporation was carried out at 2.5 kV on an Eppendorf eporator. 1 mL of THB containing 12.5% PEG 8000, 20 mM MgCl_2_, and 2 mM CaCl_2_ was immediately added and then the entire reaction was transferred to a glass culture tube. Outgrowth was at 28°C for 2 h followed by plating on THB agar supplemented with 5 µg/mL erythromycin. Plates were incubated for 48 h at 28°C. A single colony was cultured overnight in 5 mL

THB with 5 µg/mL erythromycin at 28°C. The culture was screened via PCR for the plasmid insert with the initial denaturing step extended to 10 min. The overnight culture was diluted 1:1000 THB containing 5 µg/mL erythromycin and incubated overnight at 37°C to promote single cross over events. The culture was then serial diluted and plated on THB agar plates with antibiotic and incubated at 37°C overnight. Colonies were screened for single crossover events by PCR. Single crossover colonies were inoculated in 5 mL THB at 28°C to promote double crossover events. Overnight cultures were diluted 1:1000 into 5 mL THB containing sterile 0.75 M sucrose and incubated at 37°C. Overnight cultures were serial diluted and plated on THB agar and incubated at 37°C overnight. Colonies were patched onto THB agar with and without 5 µg/mL erythromycin to confirm loss of plasmid. Colonies were also screened by PCR for the loss of *mprF*. Colonies positive for the loss of *mprF* were inoculated into 5 mL THB at 37°C. Cultures were stocked and gDNA extracted as described above, with minor modifications. Sequence confirmation of the *mprF* knockout was done via Sanger sequencing (Massachusetts General Hospital DNA Core or CU Anschutz Molecular Biology Core). The mutant was grown overnight in 15 mL THB at 37°C and pelleted at 6,150 x *g* for 5 min in a Sorvall RC6+ centrifuge at room temperature for lipid extraction as described. Genomic DNA of COH1Δ*mprF* was isolated as described above and whole genome sequencing was performed in paired-end reads (2 by 150 bp) on the Illumina NextSeq 550 platform at the Microbial Genome Sequencing Center (Pittsburgh, PA). Illumina sequence reads are deposited in the Sequence Read Archive, accession PRJNA675025.

### Complementation of *mprF* in COH1Δ*mprF* and CJB111*ΔmprF*

Electrocompetent GBS strains were generated as previously described (44). Briefly, GBSΔ*mprF* was inoculated into 5 mL THB with 0.6% glycine and grown overnight. The culture was expanded to 50 mL in pre-warmed THB with 0.6% glycine and grown to an OD_600_ nm of 0.3 and pelleted for 10 min at 3200 x *g* at 4°C in a Sorvall RC6+ floor centrifuge. The pellet was kept on ice through the remainder of the protocol. The pellet was washed twice with 25 mL and once with 10 mL of cold 0.625 M sucrose and pelleted as above. The cell pellet was resuspended in 400 µL of cold 20% glycerol, aliquoted in 50 µL aliquots, and used immediately or stored at -80°C until use. Electroporation was performed as described above, with recovery in THB supplemented with 0.25 M sucrose, and plated on THB agar with kanamycin at 300 µg/mL.

### Acidic Bligh-Dyer extractions

Centrifugation was performed using a Sorvall RC6+ centrifuge. Cultures were pelleted at 4,280 x *g* for 5 min at room temperature unless otherwise stated. The supernatants were removed, and cell pellets were stored at -80°C until acidic Bligh-Dyer lipid extractions were performed as described (3). Briefly, cell pellets were resuspended in 1X PBS (Sigma-Aldrich) and transferred to Coring Pyrex glass tubes with PTFR-lined caps (VWR), followed by 1:2 vol:vol chloroform:methanol addition. Single phase extractions were vortexed periodically and incubated at room temperature for 15 minutes before 500 x *g* centrifugation for 10 min. A two-phase Bligh-Dyer was achieved by addition of 100 µL 37% HCl, 1 mL CHCl_3_, and 900 µl of 1X PBS, which was then vortexed and centrifuged for 5 min at 500 x *g*. The lower phase was removed to a new tube and dried under nitrogen before being stored at -80°C prior to lipidomic analysis.

### Liquid Chromatography/Electrospray Ionization Mass Spectrometry

Normal phase LC was performed on an Agilent 1200 quaternary LC system equipped with an Ascentis silica HPLC column (5 µm; 25 cm by 2.1 mm; Sigma-Aldrich) as described previously (45, 46). Briefly, mobile phase A consisted of chloroform-methanol-aqueous ammonium hydroxide (800:195:5, vol/vol), mobile phase B consisted of chloroform-methanol-water-aqueous ammonium hydroxide (600:340:50:5, vol/vol), and mobile phase C consisted of chloroform-methanol-water-aqueous ammonium hydroxide (450:450:95:5, vol/vol). The elution program consisted of the following: 100% mobile phase A was held isocratically for 2 min, then linearly increased to 100% mobile phase B over 14 min, and held at 100% mobile phase B for 11 min. The LC gradient was then changed to 100% mobile phase C over 3 min, held at 100% mobile phase C for 3 min, and, finally, returned to 100% mobile phase A over 0.5 min and held at 100% mobile phase A for 5 min. The LC eluent (with a total flow rate of 300 µl/min) was introduced into the ESI source of a high-resolution TripleTOF5600 mass spectrometer (Sciex, Framingham, MA). Instrumental settings for negative-ion ESI and MS/MS analysis of lipid species were: IS = -4,500 V, CUR = 20 psi, GSI = 20 psi, DP = -55 V, and FP = -150V. Settings for positive-ion ESI and MS/MS analysis were: IS = +5,000 V, CUR = 20 psi, GSI = 20 psi, DP = +55 V, and FP = +50V. The MS/MS analysis used nitrogen as the collision gas. Data analysis was performed using Analyst TF1.5 software (Sciex, Framingham, MA).

### pH-adjusted THB growth

Approximately 30 mL of fresh THB were adjusted to different pH values, measured using a Mettler Toledo FiveEasy pH/MV meter, and sterile-filtered using 0.22 µM syringe filters. A final volume of 200 µL culture medium was aliquoted per well in a flat-bottom 96 well plate (Falcon); culture media were not supplemented with antibiotics. Overnight cultures of GBS strains were used to inoculate the wells to a starting OD_600nm_ 0.02 per well. Plates were incubated for 24 h at 37°C before OD_600nm_ was read using a BioTek MX Synergy 2 plate reader. This experiment was performed in biological triplicate.

### hCMEC cell adherence and invasion assays

Human Cerebral Microvascular Endothelial cells hCMEC/D3 (obtained from Millipore) were grown in EndoGRO-MV complete media (Millipore, SCME004) supplemented with 5% fetal bovine serum (FBS) and 1 ng/ml fibroblast growth factor-2 (FGF-2; Millipore). Cells were grown in tissue culture treated 24 well plates and 5% CO_2_ at 37°C.

Assays to determine the total number of bacteria adhered to host cells or intracellular bacteria were performed as described previously (24, 25). Briefly, bacteria were grown to mid log phase (OD_600nm_ 0.4-0.5) and normalized to 1 × 10^8^ to infect cell monolayers at a multiplicity of infection (MOI) of 1 (1 × 10^5^ CFU per well). The total cell-associated GBS were recovered after 30 min incubation. Cells were washed slowly five times with 500 µL 1X PBS (Sigma) and detached by addition of 100 µL of 0.25% trypsin-EDTA solution (Gibco) and incubation for 5 min before lysing the eukaryotic cells with the addition of 400 µL of 0.025% Triton X-100 (Sigma) and vigorous pipetting. The lysates were then serially diluted and plated on THB agar and incubated overnight to determine CFU. Bacterial invasion assays were performed as described above except infection plates were incubated for 2 h before incubation with 100 μg gentamicin (Sigma) and 5 μg penicillin (Sigma) supplemented media for an additional 2 h to kill all extracellular bacteria, prior to being trypsinized, lysed, and plated as described. Experiments were performed in biological triplicate with four technical replicates per experiment.

### Murine model of GBS hematogenous meningitis

All animal experiments were conducted under the approval of the Institutional Animal Care and Use Committee (#00316) at the University of Colorado Anschutz Medical Campus and performed using accepted veterinary standards. The murine meningitis model was performed as previously described (25, 47, 48). Briefly, 7-week-old male CD1 (Charles River) mice were challenged intravenously with 1 × 10^9^ CFU of WT COH1 or the isogenic Δ*mprF* mutant. At 72 h post-infection, mice were euthanized and blood, lung and brain tissue were harvested, homogenized, and serially diluted on THB agar plates to determine bacterial CFU.

### Histology and ELISA

Mouse brain tissue was frozen in OCT compound (Sakura) and sectioned using a CM1950 cryostat (Leica). Sections were stained using hematoxylin and eosin (Sigma) and images were taken using a BZ-X710 microscope (Keyence). Images were analyzed using ImageJ software. Meningeal thickening was quantified from sections taken from three different mice per group, and six images per slide. Meningeal thickening was quantified across two points per image. KC protein from mouse brain homogenates was detected by enzyme-linked immunosorbent assay according to the manufacturer’s instructions (R&D systems).

## Ethics Statement

Animal experiments were approved by the Institutional Animal Care and Use Committee (IACUC) at University of Colorado Anschutz Medical Campus protocol #00316 and were performed using accepted veterinary standards. The University of Colorado Anschutz Medical Campus is AAALAC accredited; and its facilities meet and adhere to the standards in the “Guide for the Care and Use of Laboratory Animals”.

## Acknowledgements

We thank Kathryn Patras at the University of California San Diego for the CNCTC 10/84 strain and Moutusee Islam in Kelli Palmer’s lab at The University of Texas at Dallas for *E. faecium* 1,231,410 DNA.

The work was supported in part by T32 5T32AI052066-18 and F31 AI164674 for H.S.M, the Coordenação de Aperfeiçoamento de Pessoal de Nível Superior, Brazil (CAPES; finance code 001 to J.D.C.M), by grants R01NS116716 and R01AI153332 from the National Institutes of Health (NIH) to K.S.D and associated NIH/NINDS supplement from R01NS116716 to R.V, NIH grant R21AI130666 and the Cecil H. and Ida Green Chair in

Systems Biology Science to K.P, NIH grant R56AI139105 to K.P and Z.G, and NIH grant U54GM069338 to Z.G.

## Supplemental Figures, and Tables

**S1 Fig.**
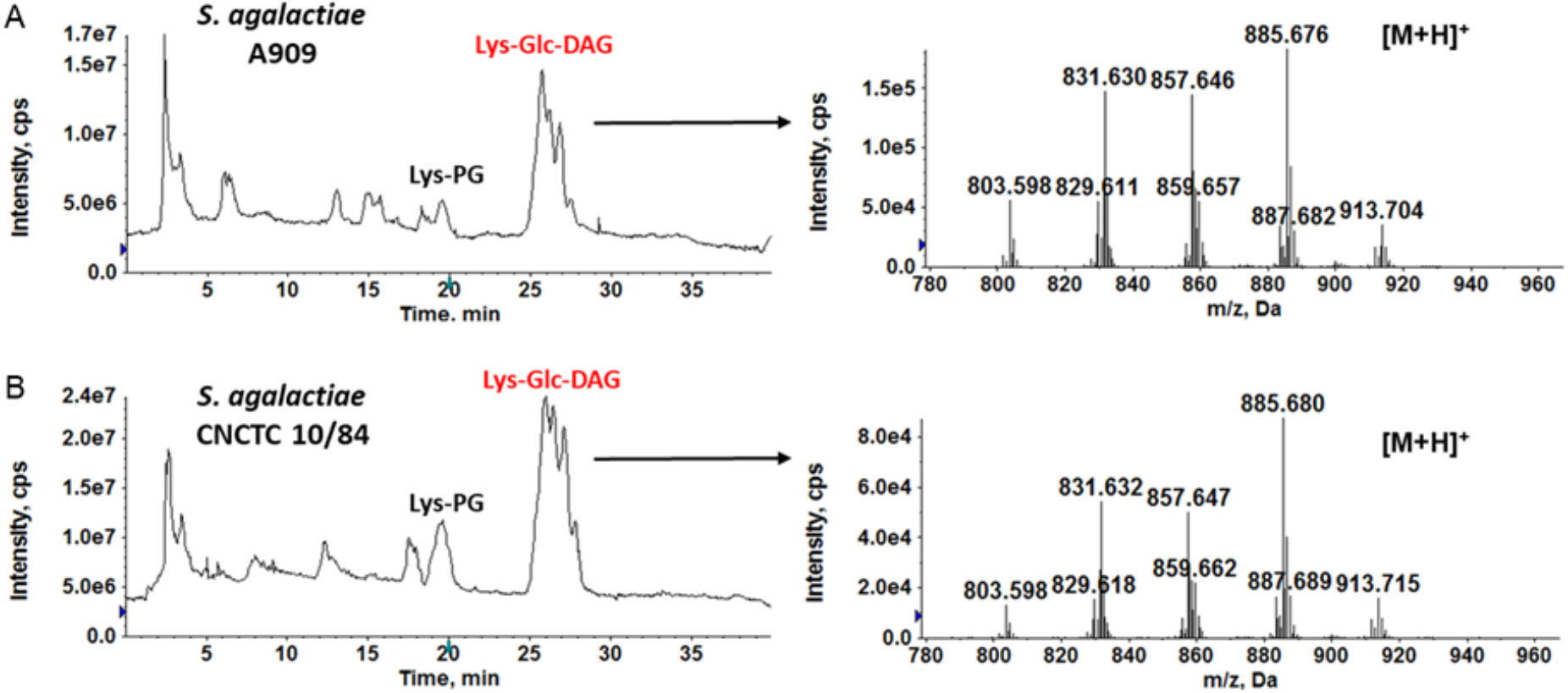
Detection of Lys-PG and Lys-Glc-DAG in *S. agalactiae* A909 and *S. agalactiae* CNCTC 10/84. Positive TICs (left panels) showing the presence of Lys-PG and Lys-Glc-DAG in S. *agalactiae* A909 and S. *agalactiae* CNCTC 10/84. Mass spectra (right panels) show the [M+H]^+^ ions of Lys-Glc-DAG.

**S2 Fig.**
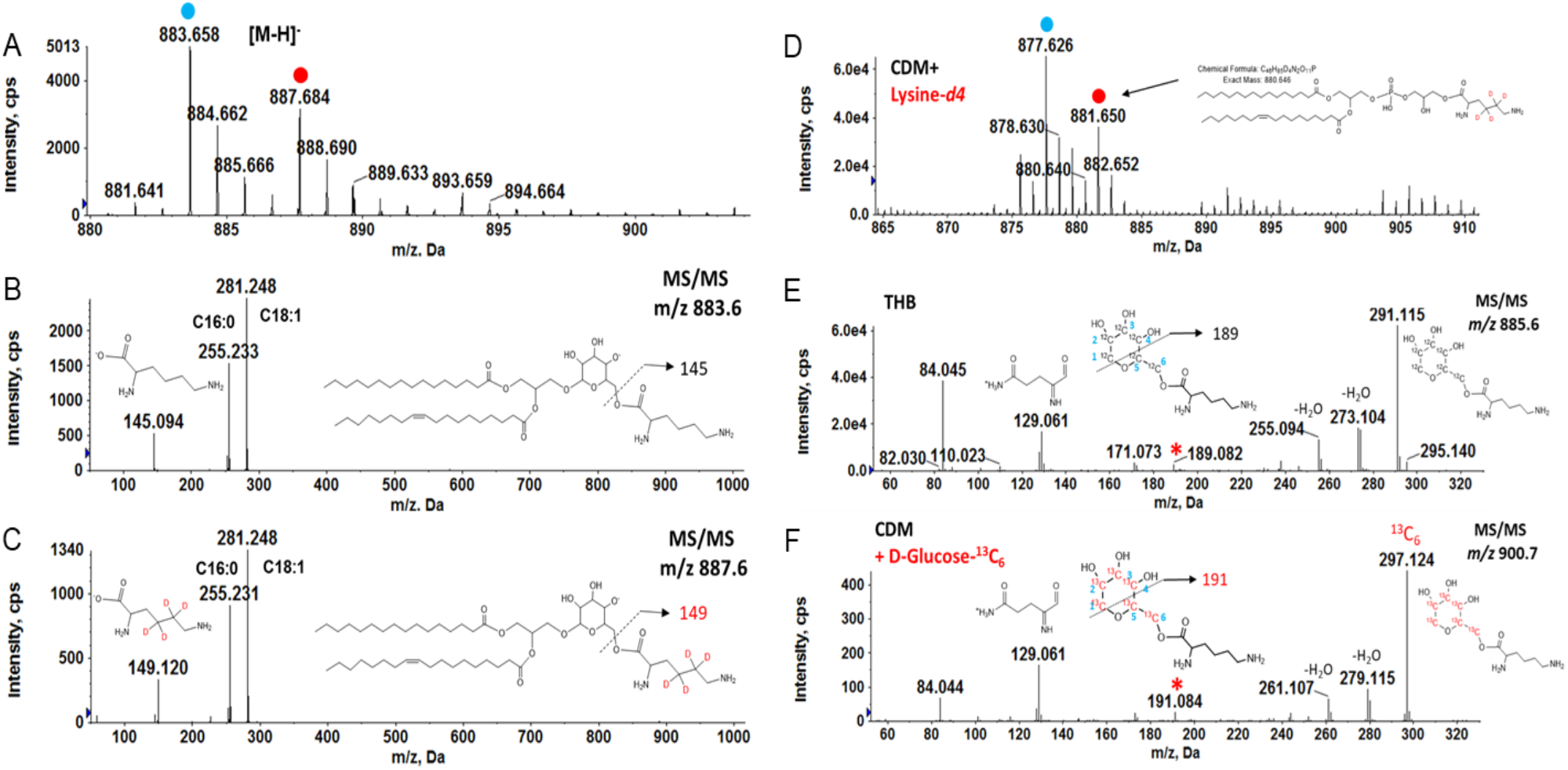
Isotopic incorporation of deuterated lysine and ^13^C-labeled glucose into Lys-Glc-DAG and Lys-PG. The lipid extracts of *S. agalactiae* COH1 cultured in DM, DM supplemented with 450 µM L-lysine-*d4* (4,4,5,5-D4), or in DM containing 0.5% w/v D-Glucose (U-^13^C_6_) were analyzed by LC-ESI/MS in the positive ion mode. A) Negative ESI/MS of [M-H]^-^ ions of major Lys-Glc-DAG species in *S. agalactiae* COH1 when cultured in DM supplemented with lysine-*d4*. The incorporation of lysine-*d4* into Lys-Glc-DAG is evidenced by an upward *m/z* shift of 4 Da of the [M-H]^-^ ion (from *m/z* 883 to *m/z* 887). B) MS/MS of [M-H]^-^ at *m/z* 883.6 produces a deprotonated lysine residue at *m/z* 145. C) MS/MS of [M-H]^-^ at *m/z* 887.6 produces a deprotonated lysine-*d4* residue at *m/z* 149. D) [M+H]^+^ ions of major Lys-PG species in *S. agalactiae* COH1 cultured in DM supplemented with lysine-*d4*. The incorporation of lysine-*d4* in Lys-PG is evidenced by an upward *m/z* shift of 4 Da from unlabeled Lys-PG (blue dot) to labeled Lys-PG (red dot). E) MS/MS of 885.6. A major product ion at *m/z* 291.1 is derived from glucose-lysine residue. F) MS/MS of *m/z* 900.7 (containing fifteen ^13^C atoms). The presence of *m/z* 297.1 (with 6 Da shift) is consistent with glucose in Lys-Glc-DAG is replaced with D-Glucose (U-^13^C_6_). The other nine ^13^C atoms are incorporated into the DAG portion of Lys-Glc-DAG. Furthermore, MS/MS data indicate that lysine is linked to the C6 position of glucose by the fragmentation schemes for forming *m/z* 189 ion from the unlabeled Lys-Glc-DAG and *m/z* 191 ion from the ^13^C-labeled Lys-Glc-DAG.

**S3 Fig.**
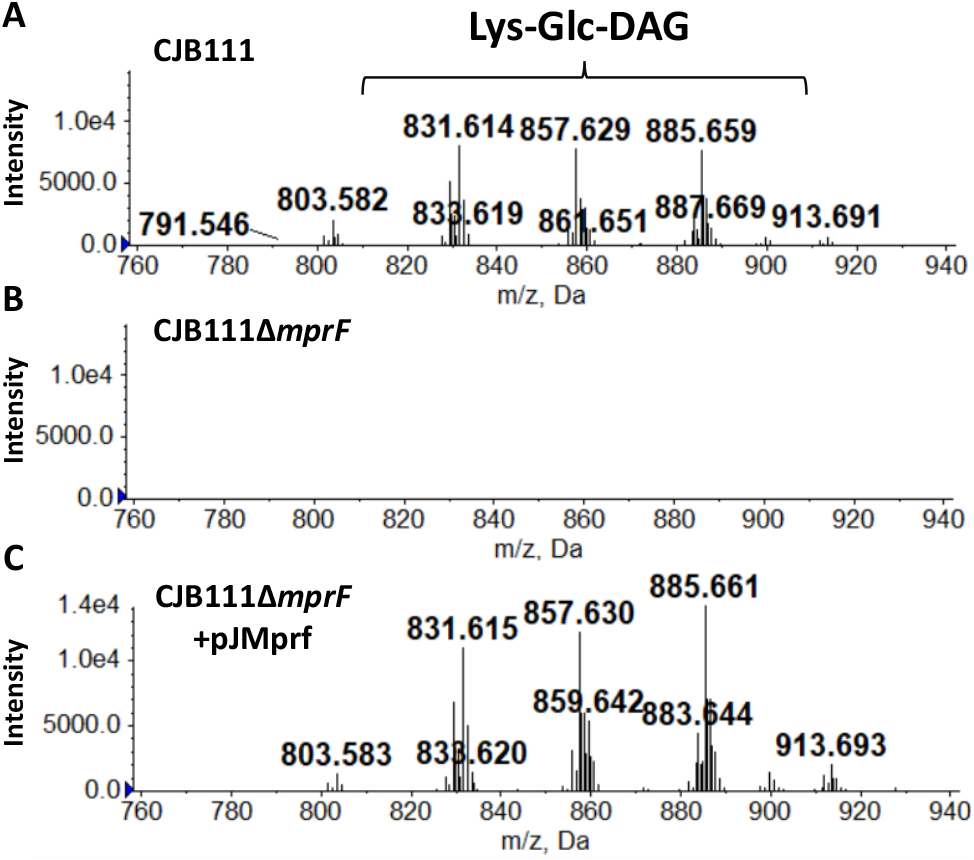
Positive ion mass spectra of retention time 27-29 minutes of hypervirulent CJB111 strain. Lys-Glc-DAG is present in the membrane of WT CJB111 (A). Deletion of *mprF* from CJB111 genome results in loss of Lys-Glc-DAG from the membrane (B). *MprF* complemented *in trans* reestablishes Lys-Glc-DAG back into the membrane (C).

**S4 Fig.**
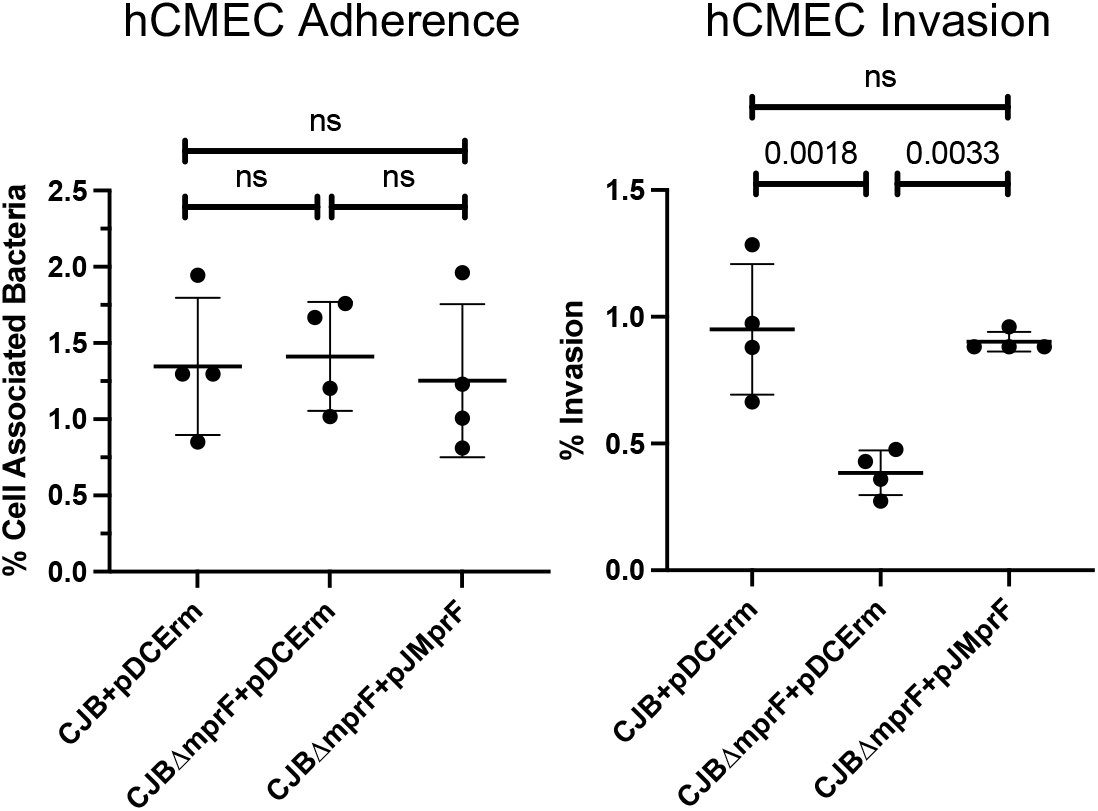
*In vitro* hCMEC adhesion and invasion of CJB111 strains. *In vitro* assays for adherence and invasion of hCMEC cells indicates *mprF* contributes to invasion but not adherence to brain endothelium. Data indicates the percentage of the initial inoculum that was recovered. Experiments were performed three times with each condition in quadruplicate. Data from one representative experiment is shown, mean and standard deviation indicated. One-Way ANOVA with Tukey’s multiple comparisons statistical test was used. P-values indicated; ns, not significant.

**S1 Table.** Observed and calculated exact masses of the [M+H]^+^ ions of Lys-Glc-DAG molecular species in *S. agalactiae* COH1. [M+H]^+^.

| Lys-Glc-DAG<sup>1</sup> | [M+H]<sup>+</sup> |  |
| --- | --- | --- |
|  | Observed mass | Exact mass |
| C28:1 | 801.575 | 801.583 |
| C28:0 | 803.595 | 803.599 |
| C30:1 | 829.610 | 829.615 |
| C30:0 | 831.624 | 831.630 |
| C32:2 | 855.623 | 855.630 |
| C32:1 | 857.644 | 857.646 |
| C34:2 | 883.657 | 883.622 |
| C34:1 | 885.672 | 885.677 |
| C36:2 | 911.686 | 911.693 |
| C36:1 | 913.703 | 913.709 |

<sup>1</sup>The numbers before and after colons indicate the total acyl chain carbon atoms and double bonds, respectively.

**S2 Table.**
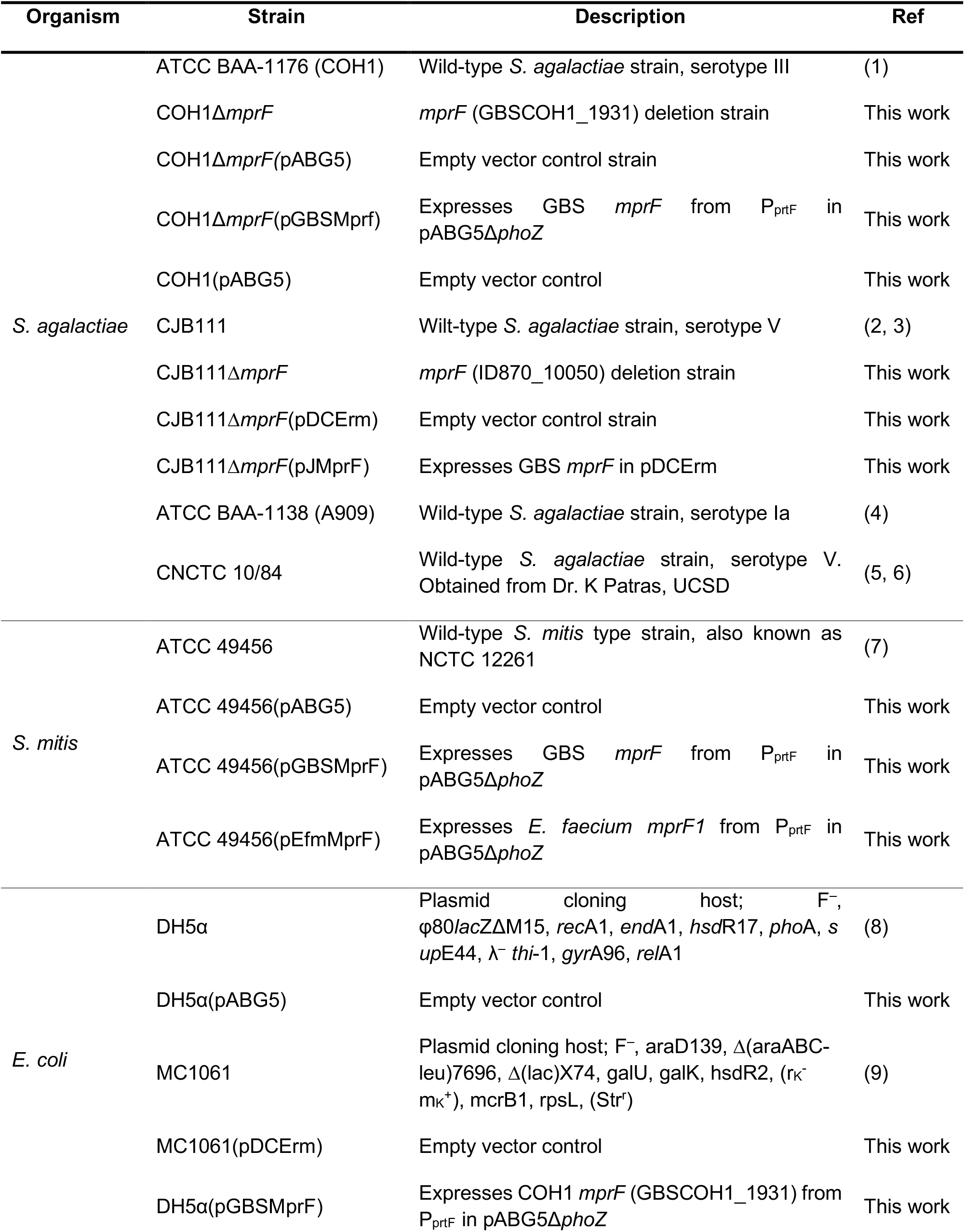
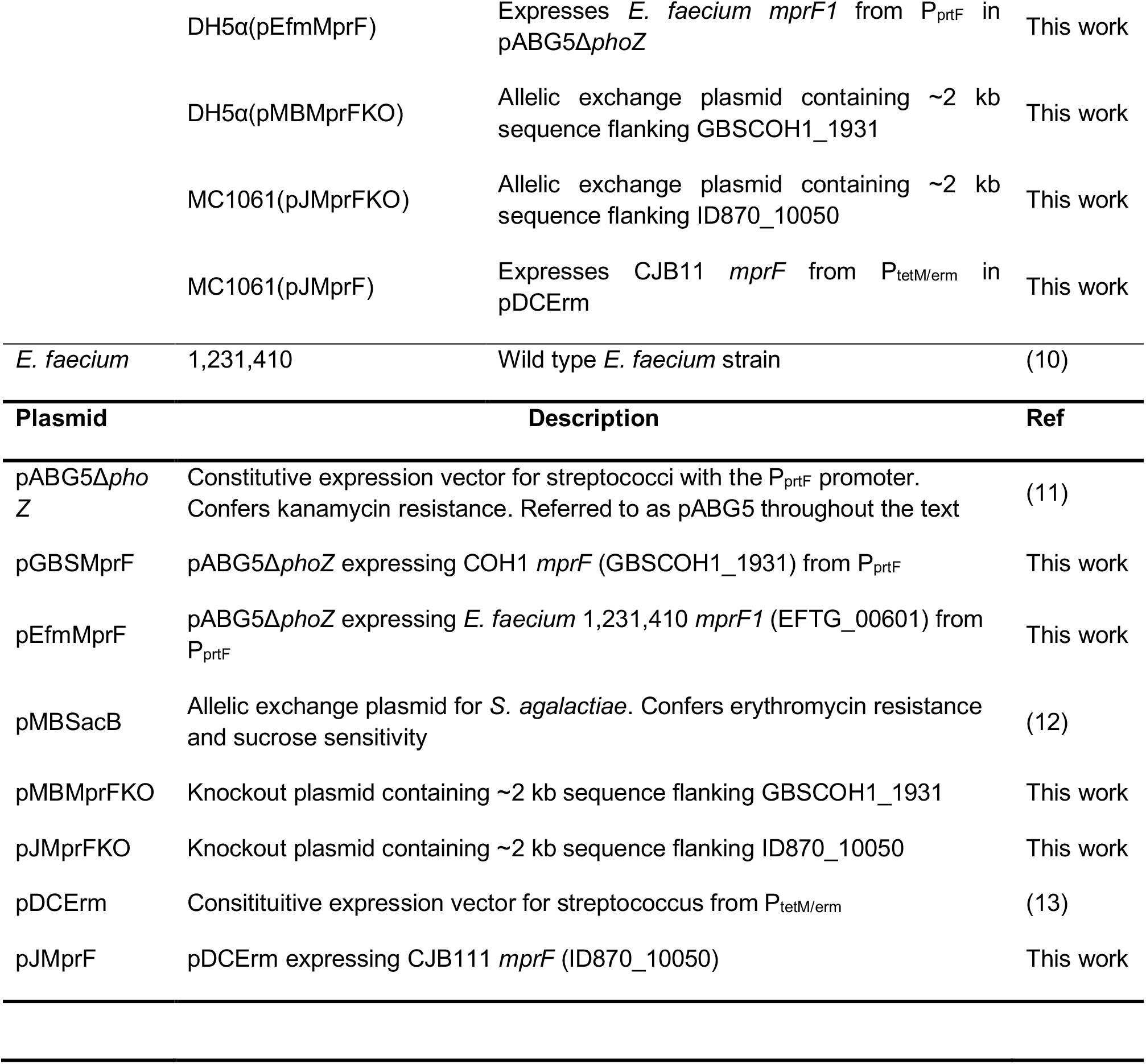
Strains and plasmids used in this study.

**S3 Table.**
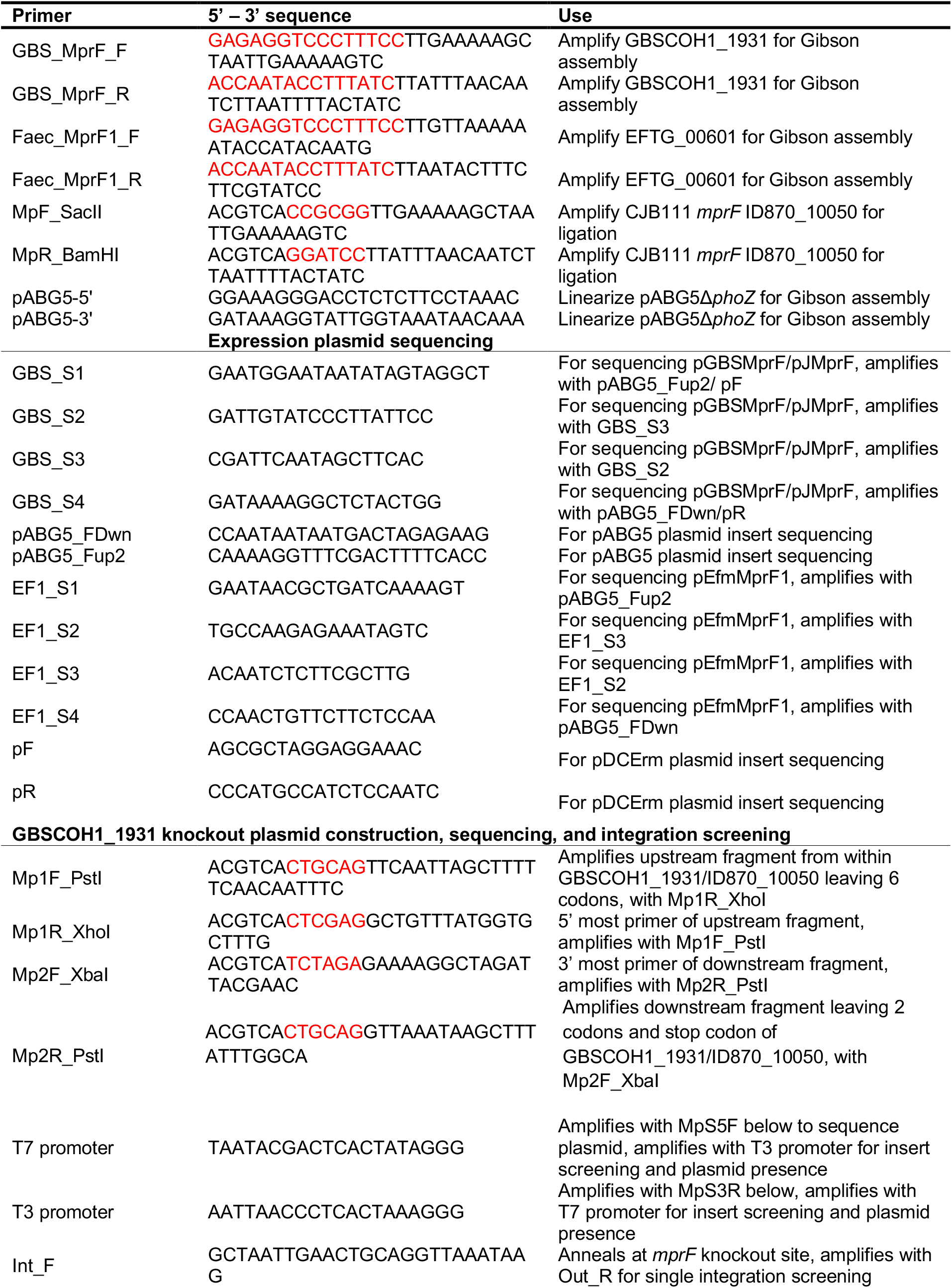
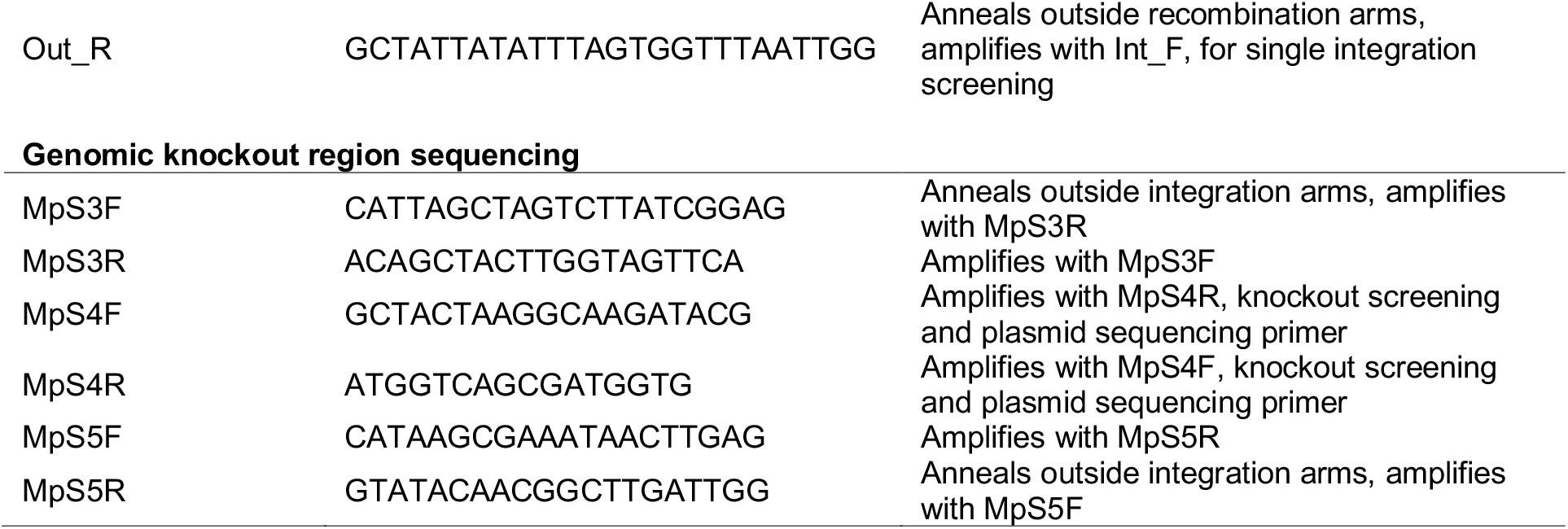
Primers used in this study.

